# Intramolecular regulation of anillin during cytokinesis

**DOI:** 10.1101/726471

**Authors:** Daniel Beaudet, Nhat Pham, Noha Skaik, Alisa Piekny

**Affiliations:** Department of Biology, Concordia University, Montreal, QC, Canada, H4B 1R6

**Keywords:** Cytokinesis, Anillin, RhoA, Importin, Ran

## Abstract

Cytokinesis occurs by the ingression of an actomyosin ring that cleaves a cell into two daughters. This process is tightly controlled to avoid aneuploidy, and we previously showed that active Ran coordinates ring positioning with chromatin. Active Ran is high around chromatin, and forms an inverse gradient to cargo-bound importins. We found that the ring component anillin contains an NLS that binds to importin and is required for its function. Anillin contains a RhoA-binding domain (RBD), which we revealed autoinhibits the adjacent NLS-containing C2 domain. Here, we show that active RhoA relieves inhibition of the C2 domain. Furthermore, FRAP experiments show that the NLS regulates anillin’s cortical properties, supporting feedback to the RBD. Indeed, mutations that disrupt the interface between the RBD and C2 domain disrupt anillin’s localization and function. Thus, active RhoA induces a conformational change that increases accessibility to the C2 domain, which is maintained by importin-binding for recruitment to the equatorial cortex.

## INTRODUCTION

Cytokinesis occurs at the end of mitosis to physically separate a cell into two daughters. This process must be strictly controlled to ensure that each daughter inherits the appropriate genetic material and cell fate determinants. Cytokinesis occurs due to the ingression of an actomyosin ring (Green et al., 2012). Contractile proteins localize broadly around the cortex in early anaphase and are restricted to the equatorial zone prior to furrow ingression. Ring assembly and ingression is regulated by active RhoA, which binds to effectors that regulate F-actin and activate non-muscle myosin II (Piekny et al., 2005, Basant and Glotzer, 2018). Active RhoA also recruits the scaffold protein anillin, which crosslinks the actomyosin ring to the plasma membrane and feeds back to stabilize active RhoA for robust cytokinesis (Piekny and Glotzer, 2008; van Oostende Triplet et al., 2014; Sun et al., 2015; Budnar et al., 2019). Since anillin crosslinks key components of the cell during cytokinesis, understanding its molecular regulation can provide insight to how cytokinesis is regulated. In particular, the C-terminus of anillin contains a RhoA-GTP binding domain (RBD), a neighboring C2 domain, and a Pleckstrin Homology (PH) domain, which coordinate its localization through 1) binding to active RhoA, 2) binding to phospholipids, importins and/or microtubules via the C2 domain, and 3) binding to phospholipids and/or septins via the PH domain (Oegema et al., 2000; Piekny and Glotzer, 2008; Piekny and Maddox, 2010; Liu et al., 2012; van Oostende Triplet et al., 2014; Sun et al., 2015; Beaudet et al., 2017). However, which of these interactions is crucial for anillin function and how they are coordinated is not clear.

Multiple mechanisms regulate cytokinesis and are derived from spindle-dependent (*e.g*. Dechant and Glotzer, 2003; Somers and Saint, 2003; Bement et al., 2005; Bringmann and Hyman, 2005; Yüce et al., 2005; Zhao and Fang, 2005; Werner et al., 2007; Lewellyn et al., 2010; van Oostende Triplet et al., 2014; Pollard and O’Shaughnessy, 2019) and -independent pathways (*e.g*. von Dassow et al., 2009; Cabernard et al., 2010; Schenk et al., 2010; Sedzinski et al., 2011; Kiyomitsu and Cheeseman, 2013; Zanin et al., 2013; Rodrigues et al., 2015; Beaudet et al., 2017). The central spindle forms between segregating chromosomes during anaphase and contains antiparallel bundled microtubules. Centralspindlin, a heterotetramer of Cyk-4 and MKLP1, bundles central spindle microtubules and forms a complex with the RhoA GEF Ect2, which stimulates the activation of RhoA in the equatorial cortex (Mishima et al., 2002; Mishima et al., 2004; Bement et al., 2005; Yüce et al., 2005; Petronczki et al., 2007; Burkard et al., 2009; Wolfe et al., 2009). Several studies showed that the membrane-localization of this complex is crucial for Ect2’s GEF activity vs. spindle localization and the role of microtubules remains to be clarified (Frenette et al., 2012; Lakomtsev et al., 2012; Basant et al., 2015; Kotýnková et al., 2016). Astral microtubules emanate toward the poles and inhibit contractility at the nearby cortex. Cells treated with nocodazole to depolymerize microtubules cause a global increase in active RhoA and an increase in the breadth of accumulated contractile proteins (Chang et al., 2008; Murthy and Wadsworth, 2008; Zanin et al., 2013; van Oostende Triplet et al., 2014). Increasing the density of astral microtubules by depleting MCAK (a microtubule depolymerase) decreases the zone of accumulated contractile proteins, which is restored upon co-depletion of anillin (Rankin and Wordeman, 2010; Zanin et al., 2013; van Oostende Triplet et al., 2014). Anillin directly binds to microtubules, and its localization to microtubules in cells is enhanced by increased microtubule stability or density, and decreased by active RhoA (Tse et al., 2011; van Oostende Triplet et al., 2014). Thus, astral microtubules may sequester anillin at the cell poles, but not at the furrow to prevent contractility at the polar cortex (Tse et al., 2011; van Oostende Triplet et al., 2014). A recent study showed that in the early *C. elegans* embryo, the clearing of F-actin and anillin at the polar cortex depends on the astral microtubule-based TPXL-1-mediated activation of Aurora A kinase (Mangal et al., 2018). However, it is unclear if Aurora A activity directly impacts RhoA and/or anillin localization or if there are other proteins involved since few other molecular regulators of the astral pathway are known (Bringmann et al., 2007; Basant and Glotzer, 2018).

Chromatin-sensing is one of the microtubule-independent mechanisms that regulates cytokinesis. Importins bind to the nuclear localization signals (NLS) on cargo to mediate their nuclear transport (Soniat and Chook, 2015). Active Ran is enriched in the nucleus and competes with importins to release cargo. During mitosis, the Ran gradient persists around chromatin where it controls spindle assembly (Li and Zheng, 2004; Kaláb et al., 2006; Clarke and Zhang, 2008; Kaláb and Heald, 2008). The dogma is that importin-binding to the NLS of a spindle regulator sterically blocks interactions with partners required for their function (Weaver and Walczak, 2015). Thus, active complexes assemble the spindle near chromatin where active Ran is high. However, findings from our lab and others supports a very different function for the regulation of cortical proteins by importin-binding (Deng et al., 2007; Dehapiot et al., 2013; Dehapiot and Halet, 2013; Samwer et al., 2013; Kiyomitsu and Cheeseman, 2013; Beaudet et al., 2017). For example, importins function as a ruler during meiosis or cytokinesis, where optimal levels facilitate polarization of the nearby cortex (Deng et al., 2007; Dehapiot et al., 2013; Dehapiot and Halet, 2013; Samwer et al., 2013; Beaudet et al., 2017). This mechanism senses chromatin position to ensure that the polar body or contractile ring are properly positioned. A recent study showed that importin-a is lipid-modified by palmitoylation in *Xenopus* egg extracts, and the authors proposed this creates a cortical sink of importin-a to reduce cytoplasmic levels for scaling spindle and nuclear size (Brownlee and Heald, 2019). It will be interesting to determine if palymitoylated importin-a can form functional complexes and mediate processes at the membrane.

Previous studies in our lab provided mechanistic insight to how importins regulate anillin. We found the NLS of anillin is required for its localization and function during cytokinesis (Beaudet et al., 2017). In addition, anillin contains a RhoA-binding domain (RBD) required for cortical recruitment, and we found that the RBD autoinhibits the adjacent NLS-containing C2 domain. Our model is that binding to active RhoA facilitates a conformational change in anillin that increases accessibility to the C2 domain, and importin-binding stabilizes this conformation for optimal recruitment to the equatorial cortex. Here, we present data supporting this model. We found that active RhoA facilitates importin-binding, while inactive RhoA or mutating the RBD decreases importin-binding. Through live-imaging and FRAP experiments we found that mutating the NLS alters anillin’s cortical properties. Anillin’s localization and function are abolished when NLS mutations are combined with mutations that weaken the interface between the RBD and C2 domain. When stronger interface mutations are introduced into anillin, it similarly fails to localize to the furrow and fails to undergo cytokinesis. This data shows that the interface is crucial to drive feedback between the C2 domain and the RBD for cortical recruitment.

## RESULTS

### The C-terminus has different cortical retention properties compared to full-length anillin

Anillin has multiple binding domains that crosslink the main components of the cytokinesis machinery (Figure 1A). The C-terminus contains a RhoA-binding domain (RBD) that binds to active RhoA, a C2 domain with an NLS and binding sites for phospholipids, microtubules and Ect2, and a PH domain that binds to phospholipids and septins (Piekny and Maddox, 2010; Sun et al., 2015). Prior studies showed that anillin requires active RhoA for its recruitment to the equatorial cortex, and the C-terminus colocalizes with RhoA during cytokinesis. Thus, the C-terminus of anillin is now being used as a reporter for active RhoA (Piekny and Glotzer, 2008; Sun et al., 2015; Wagner and Glotzer, 2016). However, we do not fully understand how interactions in the C-terminus are coordinated for anillin’s localization and function.

**Figure 1.**
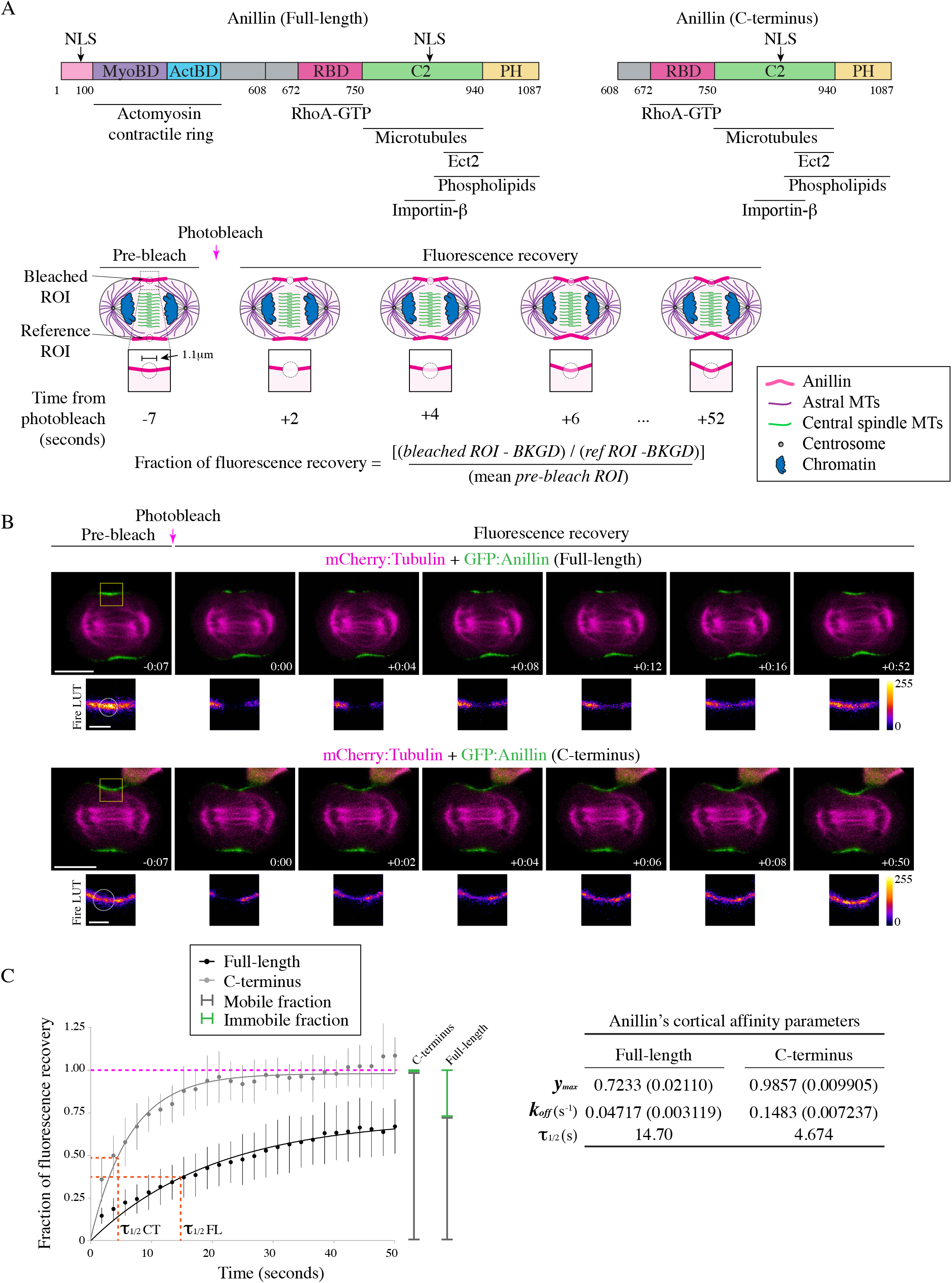
The C-terminus has different cortical properties compared to full-length anillin. A) The structures of full-length and the C-terminus of anillin are shown, with binding domains as indicated (pink = contains N-terminal NLS; purple = MyoBD, binds myosin; blue = ActBD binds actin; dark pink = RhoA-GTP binding domain (RBD), binds active RhoA; green = C2, binds microtubules, phospholipids, Ect2, and contains C-terminal NLS that binds Importin-β; yellow = Pleckstrin Homology (PH), binds phospholipids). Cartoon schematics show the fluorescence recovery after photobleaching (FRAP) of a region of interest (ROI) compared to a reference ROI in a cell expressing anillin (pink) during anaphase and early ingression. The photobleached region is shown in the boxed inset. Other components of the cell are indicated in the legend (astral microtubules in purple, central spindle microtubules in green, centrosomes in grey, and chromatin in blue). The equation used to determine the fraction of fluorescence recovery is indicated (background; BKGD). B) Timelapse images show FRAP of HeLa cells expressing mCherry:Tubulin (magenta) and GFP-tagged full-length (green; top panels) or the C-terminus of anillin (green; bottom panels). The boxed insets show the ROI’s that were photobleached in Fire LUTs. The scale bars are 10 μm or 2 μm for boxed insets. Indicated times are before (−) or after (+) photobleaching. C) A graph shows the fraction of fluorescence recovery over time for full-length (n=31) and the C-terminus (n=15) of anillin. The Y-axis shows the fraction of recovery (corrected) and the X-axis shows the time in seconds. The green lines indicate the immobile fraction, while the grey lines show the mobile fraction. Bars show standard deviation (SD). The table shows the maximum recovery (*y_max_*), dissociation rate (*k_off_*) and half-life (τ_1/2_) of full-length and C-terminus of anillin. Standard errors are shown in parentheses (SEM).

To characterize the cortical properties of anillin, we performed fluorescence recovery after photobleaching (FRAP) experiments on HeLa cells expressing GFP-tagged full-length anillin and co-expressing mCherry:Tubulin using equatorially positioned cortical regions of interest (ROI’s). The % of fluorescence recovery was determined by measuring the fluorescence signal of the photobleached ROI relative to the pre-bleached ROI over 50 seconds at 2 second intervals. The bleached ROI’s were corrected for background and photobleaching due to image acquisition by internally controlling each experiment with a reference ROI on the opposite side of the cortex (Figure 1A). First, we compared anillin’s cortical retention during early vs. late ingression (Figure S1). The stage of ingression was determined by the ratio of the length of the ingressed cortex over the width of the cell at the equator; where *R* > 0.6 was ‘early’ and *R* < 0.6 was ‘late’ (Figure S1A). There were no significant differences in the dissociation rates (k_off_) between early and late ingression, however, there was an increase in the immobile fraction during late ingression (Figure S1B and S1C). This suggests that anillin has stronger cortical retention during late cytokinesis, which may be important for the contractile ring – midbody transition.

Next, we determined how cortical properties differ between full-length anillin and the C-terminus (Figure 1B). As shown in the table, full-length anillin has greater cortical retention and a slower dissociation rate compared to the C-terminus (Figure 1C). The average k_off_ for full-length anillin was 0.047s^−1^ and recovered to a maximum of 72.33% of the pre-bleached intensity compared to C-terminus, which had an average k_off_ of 0.1483s^−1^ and recovered to 98.57% of the pre-bleached intensity (Figure 1C). The faster dissociation rate and higher recovery suggests that the C-terminus is more mobile and has lower cortical retention compared to full-length anillin.

### The C-terminal NLS regulates anillin’s cortical affinity during cytokinesis

We previously showed that the NLS influences anillin’s localization to the cortex prior to ingression, and we wanted to determine how importin-binding regulates anillin’s cortical properties (Beaudet et al., 2017; Figure 4A and 5A). First, we show that anillin directly binds to importin, since GST:Importin-β binds to MBP:Anillin (C2 domain) *in vitro* and mutating the NLS significantly reduces binding (Figure 2A). To elucidate how the loss of importin-β binding alters anillin’s turnover and retention at the equatorial cortex, we performed FRAP experiments as described for Figure 1. HeLa cells co-expressing mCherry:Tubulin and GFP-tagged C-terminus of anillin containing the NLS mutations were measured during early ingression (Figure 2B). Mutating the NLS caused a decrease in the mobile fraction and a faster dissociation rate compared to the control. For example, the NLS mutant had an average k_off_ of 0.3230s^−1^ and recovered to 73.60% of the pre-bleached signal intensity compared to control, which had an average k_off_ of 0.1483s^−1^ and recovered to 98.57% of the pre-bleached signal intensity (Figure 2B and C). This data suggests that importin-binding improves anillin’s retention by reducing dissociation, but also makes anillin more dynamic in its ability to exchange with the cortex.

**Figure 2.**
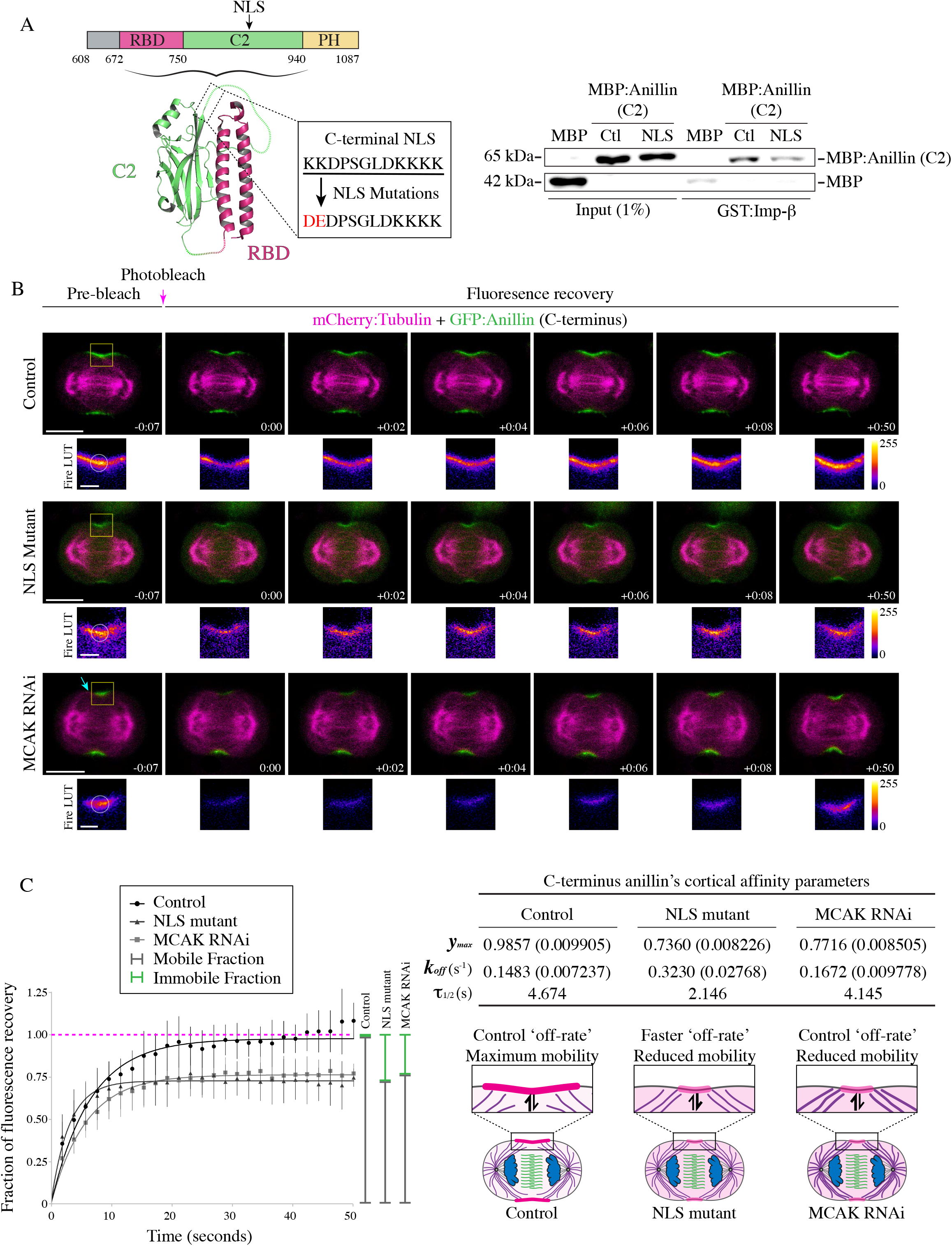
The C-terminal NLS regulates anillin’s cortical affinity during cytokinesis. A) A cartoon schematic shows the C-terminus of anillin, with binding domains and the NLS as indicated (dark pink = RBD; green = C2; yellow = PH). Underneath is a ribbon structure of the RBD (pink) and C2 domain (green). The amino acids of the NLS and those that were mutated (850 KK 851 – DE) are shown in the box. On the right, immunoblots show the *in vitro* binding of GST-tagged Importin-β (Imp-β) with MBP-tagged C2 domain (Ctl) vs. the C2 domain with the NLS mutations (NLS). Inputs are shown on the left, and pull downs on the right. B) Timelapse images show FRAP of HeLa cells expressing mCherry:Tubulin (magenta) and GFP-tagged C-terminus of anillin (green), or with the NLS mutations, or after treatment with MCAK RNAi. The boxed insets show the ROI’s that were photobleached in Fire LUTs. The scale bars are 10 μm or 2 μm for boxed insets. The blue arrow points to overextended microtubules. Indicated times are before (−) or after (+) photobleaching. C) A graph shows the fraction of fluorescence recovery (Y-axis) over time (X-axis; seconds) for control (n=15), NLS mutant anillin (n=12), or after MCAK RNAi (n=8). The green lines indicate the immobile fraction, while the grey lines show the mobile fraction. Bars show SD. The table shows the maximum recovery (*y_max_*), dissociation rate (*k_off_*) and half-life (τ_1/2_) for each condition as indicated. SEM are shown in parentheses. Below the table, cartoon schematics of cells show how mutating the NLS causes a faster off-rate vs. MCAK RNAi, which has an off-rate that is similar to control. However, both the NLS mutant and MCAK RNAi have reduced mobility.

Astral microtubules also regulate the localization of anillin during cytokinesis. We previously showed that overextending the astral microtubules with MCAK RNAi causes a delay in anillin’s cortical recruitment and localization to a narrow region of the equatorial cortex similar to the NLS mutant (Figure 2B; van Oostende Triplet et al., 2014). We also found that disrupting the astral microtubules caused NLS mutant anillin to spread along the equatorial cortex (Beaudet et al., 2017). Next, we determined how microtubules influence anillin’s cortical turnover and retention. FRAP experiments were performed using HeLa cells treated with MCAK RNAi, coexpressing mCherry:Tubulin and GFP-tagged C-terminus of anillin as described for Figure 1 and 2A (Figure 2B and C). MCAK-depleted cells had an average k_off_ of 0.1672s^−1^ and the fluorescence signal recovered to 77.16% of the pre-bleached signal. Thus, astral microtubules decrease the mobility of cortical anillin, but have no impact on its dissociation. This suggests that the NLS mutant alters a cortical property of anillin independent of microtubules.

We previously found that the microtubule-binding region of anillin maps to the C2 domain, but the precise binding site was not determined (van Oostende Triplet et al., 2014; Beaudet et al., 2017). We wanted to determine if the importin and microtubule binding regions overlap, and if their binding is competitive. First, we performed co-sedimentation assays to test how mutating the NLS changes anillin’s affinity for microtubules. This was done using 1 μM of recombinant MBP-tagged C2 domain of control or NLS mutant anillin with varying concentrations of polymerized Taxol-stabilized microtubules (0.5 – 4 μM) and quantifying the amount of protein in the pellets vs. supernatants on a Coomassie-stained gel (Figure S2A). Anillin’s affinity for microtubules was quantified by comparing bound (pellet) vs. unbound (supernatant) protein, and was found to be 0.089 μM for control anillin compared to 0.96 μM for NLS mutant anillin (Figure S2A). This ~10-fold decrease in affinity suggests that the NLS is partially required for microtubule binding. Next, we determined if importin competes with microtubules for anillin-binding. To test this, we performed co-sedimentation assays using a fixed 1:2.5 μM ratio of anillin (MBP:Anillin C2) and microtubules, and titrated in varying concentrations of recombinant GST:Importin-β (0.2 – 1.5 μM; Figure S2B). Increasing the concentration of importin-β reduced, but did not eliminate anillin-binding to microtubules (Figure S2B). This data suggests that importin and microtubules partially compete for anillin binding at a site that includes the NLS. Thus, when the NLS is mutated, cytokinesis phenotypes could be caused by a reduction in importin- and/or microtubule-binding. However, we previously showed that depolymerizing microtubules using a low dose of nocodazole increases anillin’s cortical enrichment, suggesting that reducing anillin’s affinity for microtubules would increase anillin’s retention or mobility. Since the phenotypes caused by the NLS mutant are the opposite, they more likely arise due to loss of importin-binding and a decrease in cortical affinity.

Previous studies showed that the C2 domain can bind to PI4,5P_2_ phospholipids (Sun et al., 2015; Budnar et al., 2019). Although the NLS mutations are outside of the putative lipid-binding site, we wanted to be sure that they did not impact lipid-binding. We compared the binding profiles of recombinant control GST:Anillin RBD + C2 with the NLS mutant using strips blotted with a variety of phospholipids including mono, di and tri-phosphorylated (3,4,5) lipids and found no change in their profiles (Figure S2C). Interestingly, there was a strong lipid preference for PI3P. We previously showed that GFP:Anillin C-terminus bound preferentially to PI4,5P_2_ lipids, suggesting that either the PH domain, and/or complexes formed in cells influence anillin’s lipid preference (Frenette et al., 2012).

### The NLS is auto-inhibited by the RBD and is relieved via RhoA binding

Our previous study showed that the NLS in the C2 domain is intramolecularly inhibited by the RBD, since removing the RBD increases anillin’s affinity for importin-β from cell lysates (Beaudet et al., 2017). This inhibition is direct because MBP:Anillin C2 binds more strongly to GST:Importin-β *in vitro* compared to MBP:Anillin C-terminus (Figure 3A). To determine if active RhoA relieves intramolecular inhibition from the RBD, pull down assays were performed using lysates from cells where RhoA activity was manipulated. Lysates collected from cells expressing GFP:Anillin (C-terminus) were loaded with 5 mM of GDP or GTP to generate higher levels of GTPases bound to GDP or GTP as described previously (Piekny and Glotzer, 2008; Figure 3B).

**Figure 3.**
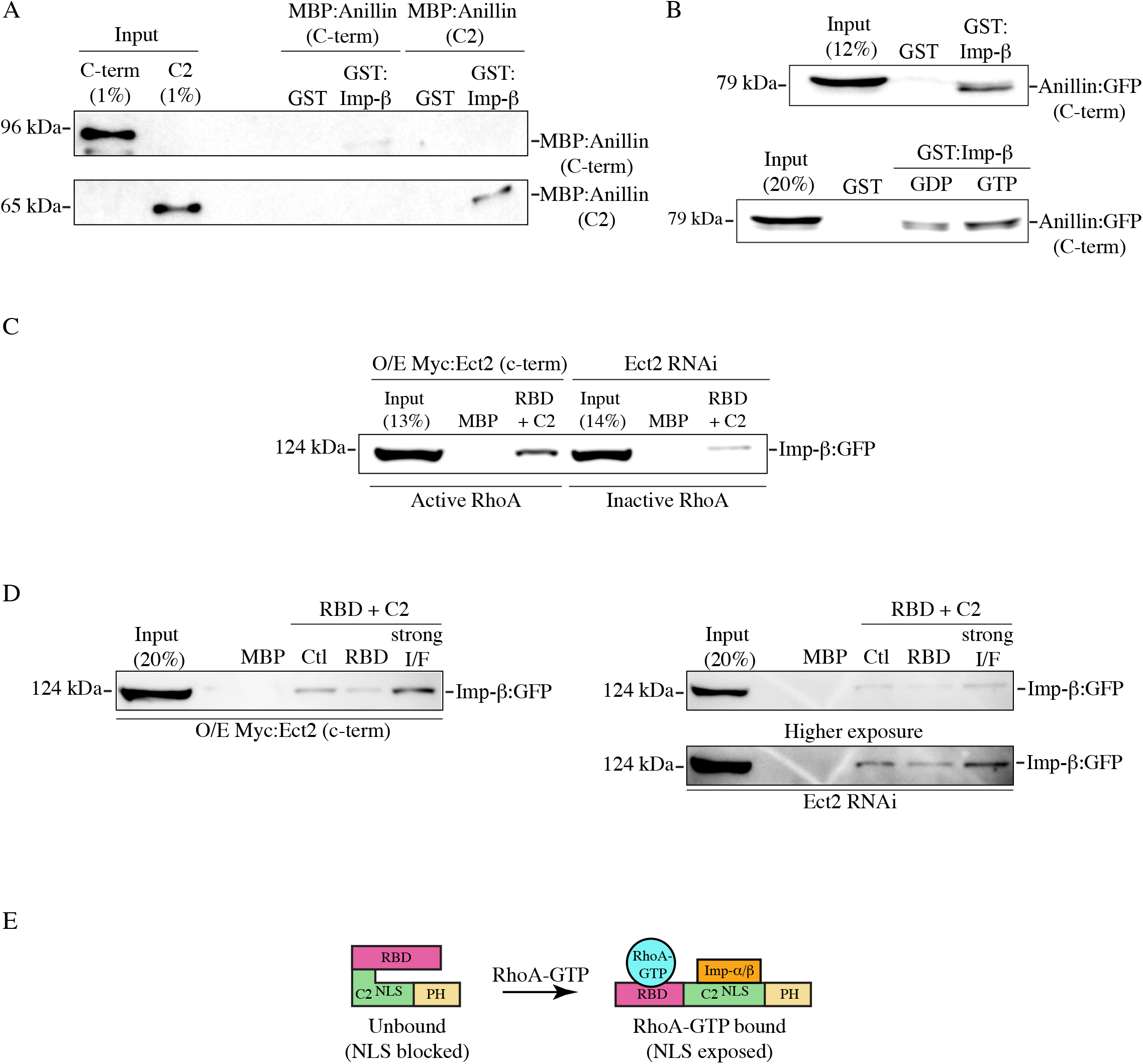
The NLS is inhibited by the RBD and is relieved via RhoA-bindings. A) Immunoblots show *in vitro* binding of purified GST or GST-tagged importin-β (Imp-β) with MBP-tagged C-terminus of anillin (C-term; top) or the C2 domain (bottom). B) Immunoblots show pull downs of GFP-tagged C-terminus of anillin (C-term) from HeLa cell lysates with purified GST or GST-tagged importin-β (top). Cell lysates were preloaded with 5 mM GDP or 5 mM GTP as indicated (bottom). C) An immunoblot shows pull downs of GFP-tagged importin-β from lysates from cells overexpressing (O/E) Myc-tagged Ect2 C-terminus to generate active RhoA (left), or after Ect2 RNAi to reduce active RhoA (right), with purified MBP or MBP-tagged anillin (RBD + C2). D) Immunoblots show pull downs of GFP-tagged importin-β from lysates as in C) with MBP, MBP-tagged anillin (RBD + C2; Ctl) or with RBD mutations (RBD), or mutations that strongly disrupt the RBD-C2 interface (strong I/F). E) A cartoon schematic shows how RhoA-GTP (blue) binding to the RBD (pink) could relieve inhibition of the C2 domain (green; NLS is indicated) to enable importin-α/β (orange) binding to the NLS (PH domain is in yellow).

GST:Importin-β pulled down more anillin from lysate loaded with GTP compared to lysate loaded with GDP. Since this assay was not selective for RhoA, we determined how altering RhoA activity impacts importin-binding to anillin. MBP:Anillin (RBD + C2) was used to pull down GFP-tagged Importin-β from lysates where the C-terminus of Ect2, which has unregulated GEF activity toward RhoA, was over-expressed in cells (Solski et al., 2004; Frenette et al., 2012), or where Ect2 RNAi was used to deplete endogenous Ect2 to reduce active RhoA (Figure 3C). We found that importin-β was more strongly pulled down with increased levels of active RhoA compared to lysates with reduced levels of active RhoA. We also determined how introducing mutations into the RBD that disrupt RhoA-binding (A703E, E721A; Sun et al., 2015) affect anillin’s affinity for importin. When MBP:Anillin (RBD + C2) containing these RBD mutations were used to pull down GFP-tagged Importin-β from lysates, importin-binding was drastically reduced compared to control (Figure 3D) regardless of Ect2 levels or activity. This data suggests that importin-binding to anillin is regulated by active RhoA, and our model is that RhoA-binding causes a conformational change that exposes the NLS in the C2 domain (Figure 3E).

### Importin-binding enhances anillin’s affinity for the cortex

Mutating the NLS reduces anillin’s cortical affinity, implying that this mutant could impact anillin’s recruitment by active RhoA. We wanted to determine how the RBD and C2 domains are coordinated to drive anillin’s localization and function for cytokinesis. To test this, we performed localization studies in HeLa cells co-expressing H2B:mRuby to visualize chromatin and GFP-tagged C-terminal control or mutant anillin constructs. Prior studies showed that decreasing active RhoA or mutating the RBD (A703E, E721A) prevents anillin’s accumulation at the equatorial cortex (Piekny and Glotzer, 2008; Sun et al., 2015). We determined how mildly disrupting the interface (I/F) between the RBD and C2 domain (843-DFEINIE to AFAINIA; Piekny and Glotzer, 2008), mutating the NLS (NLS mutant; Beaudet et al., 2017) or combining the I/F and NLS mutations affects anillin localization compared to the RBD mutant (Figure 4A). At anaphase onset, the C-terminus of anillin was broadly cortically localized, followed by its restriction to the equatorial cortex prior to ingression (Figure 4A and B). As expected, anillin failed to localize to the cortex when the RBD was mutated (Figure 4A and B). Mutating the I/F or NLS reduced the breadth of anillin’s cortical localization, causing it to be restricted to a narrow region compared to control anillin (Figure 4A and B). Interestingly, anillin failed to localize to the cortex altogether when the NLS and I/F mutations were combined, similar to the RBD mutant (Figure 4A and B). We quantified the changes in localization for the different mutants in two ways. First, we measured the breadth of anillin by drawing a line scan around the perimeter of the cell, then added the number of pixels above 50% maximum levels and determined their ratio vs. the total number of the perimeter. The ratio of anillin containing the NLS or I/F mutations was significantly lower compared to control anillin (Figure 4B). Next, we measured the ratio of cortical vs. cytosolic anillin, and found that while the I/F mutant was moderately enriched in the cytosol, the I/F + NLS mutant were strongly enriched in the cytosol similar to the RBD mutant (Figure 4C). This data suggests that RhoA and importin-binding work together to control anillin’s cortical localization during cytokinesis.

**Figure 4.**
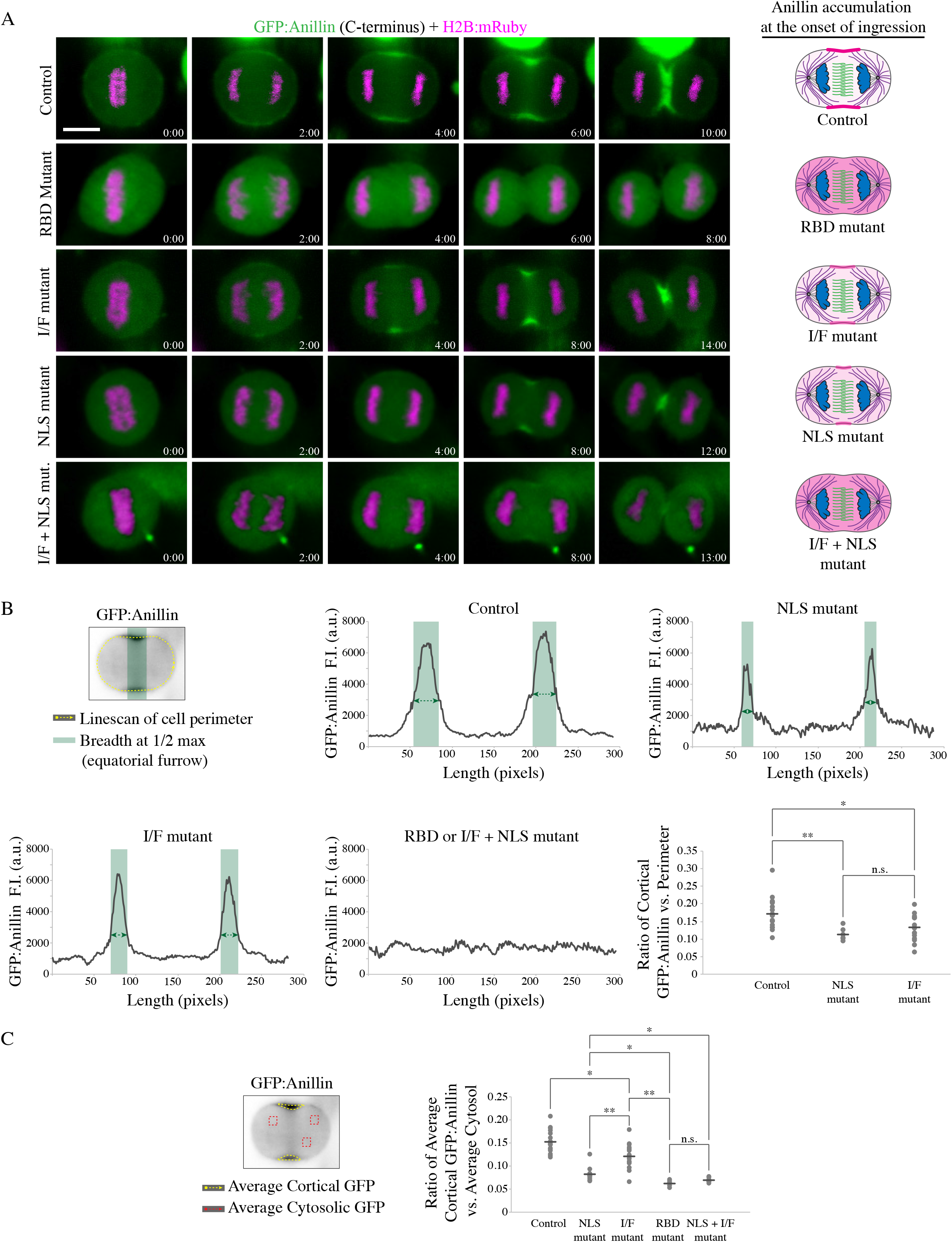
Importin-binding enhances anillin’s affinity for the equatorial cortex. A) Timelapse images show HeLa cells expressing GFP-tagged C-terminus anillin constructs (green) and H2B:mRuby (magenta; to show chromatin) as indicated: control (n=18), RhoA-binding domain mutant (RBD mutant; A703E, E721A; n=5), RBD-C2 interface mutant (I/F mutant; 843-DFEINIE to AFAINIA; n=15), NLS mutant (n=12), and a combination of the I/F + NLS mutations (I/F + NLS mutant; n=8). Times are indicated from anaphase onset. The scale bar is 10 *μ*m. Cartoon cells show changes in the distribution of anillin (pink) at the onset of ingression for the different constructs as indicated. B) Graphs of line scans are shown for the different constructs as in A), with fluorescence intensity on the Y-axis and length (pixels) on the X-axis. The line scan (dotted yellow) was drawn as shown on the cell image (upper left). Regions of the cortex above 50% maximum levels are highlighted in green and represent accumulated anillin. To the right, a dot plot shows changes in the ratio of the breadth of accumulated anillin vs. total cell perimeter (Y-axis) for the control and mutants as indicated. The means are indicated by the dark grey lines, and significance was determined using the Student’s t test (* p < 0.05, ** p < 0.001). C) The cell image (left) shows the area (yellow dotted line) measured to determine the average intensity of GFP-tagged anillin at the cortex, and the ROI’s (red boxes) used to calculate the average intensity of anillin in the cytosol. The dot plot shows the average ratio of cortical anillin vs. cytosol (Y-axis) for the control and mutants in A) as indicated. The means are indicated by the dark grey lines, and significance was determined using the Student’s t test (* p < 0.05, ** p < 0.001).

We further assessed the impact of these mutations on anillin’s function in the context of the full-length protein and its requirement for cytokinesis. Constructs containing RNAi-resistant GFP-tagged full-length anillin, or with mutations in the NLS, I/F, or both, were expressed in HeLa cells depleted of endogenous anillin and co-expressing mCherry:Tubulin to visualize the mitotic spindle. Cells were imaged after anaphase onset to assess localization and cytokinesis phenotypes (Figure 5A). As expected, anillin localized to the cortex ~2 minutes after anaphase onset and was restricted to the equatorial cortex as the contractile ring ingressed (100% of cells successfully ingressed; Figure 5A and B). Cortical recruitment of the I/F mutant was delayed until ~4 minutes after anaphase onset, and was restricted to a narrow region. A small proportion of cells expressing the I/F mutant failed cytokinesis (22.7%; Figure 5B) and those that completed ingression took longer compared to control cells (17.8 ± 4.3 mins vs. 12.3 ± 2.7 mins; Figure 5C). Cortical recruitment of the NLS mutant was also delayed and restricted to a narrow region. As previously reported, a subset of cells expressing the NLS mutant failed cytokinesis (36.4%; Figure 5A and B), and those cells that completed ingression took longer compared to control cells (16.4 ± 3.5 mins; Figure 5C). Anillin containing both the NLS and I/F mutations failed to localize to the cortex, and the majority of cells failed cytokinesis (70.6%; Figure 5A and B). Those cells that ingressed were significantly delayed compared to control cells (19.2 ± 1.6 mins; Figure 5C).

**Figure 5.**
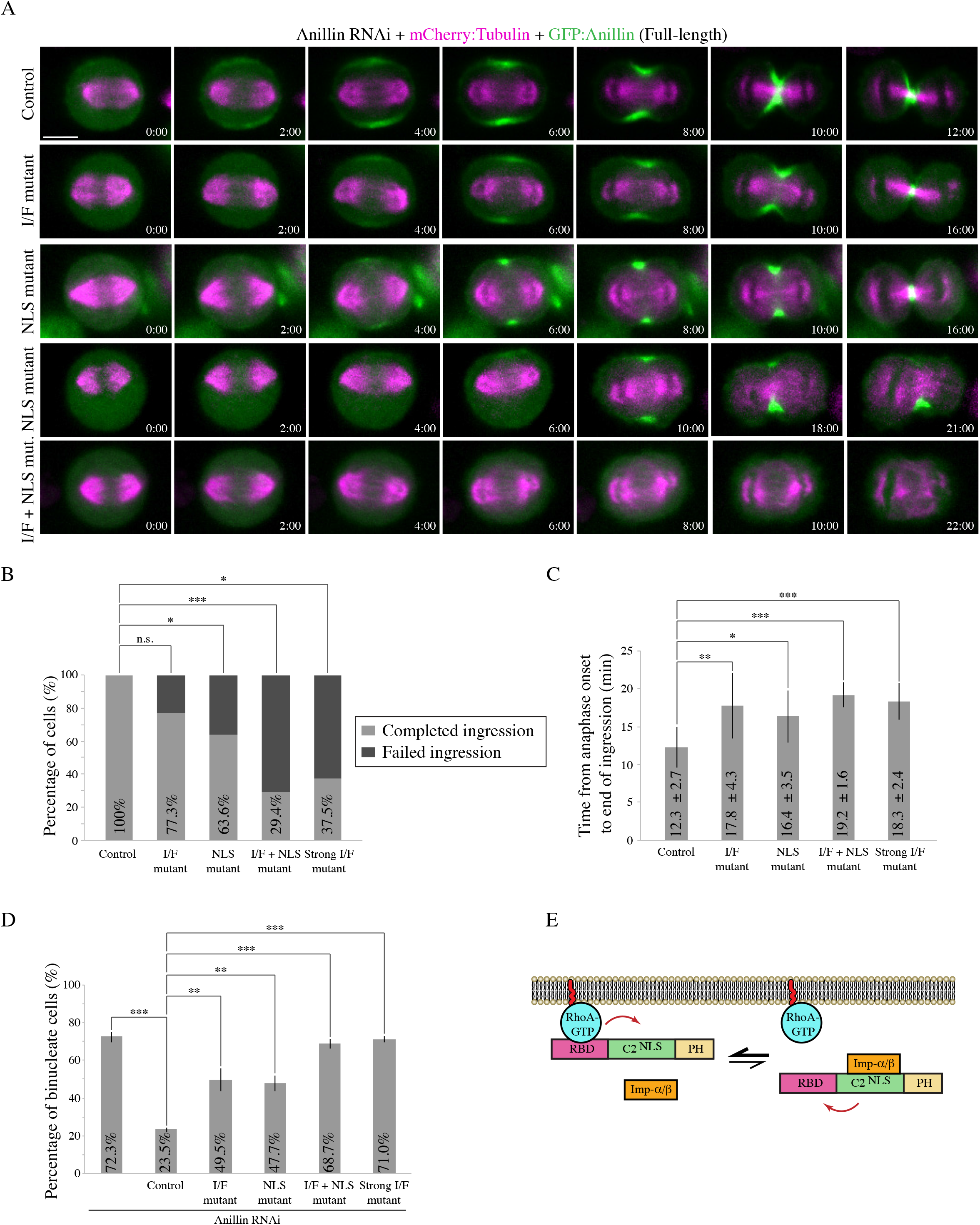
Anillin requires importin-binding and the interface between the RBD and C2 domains for cytokinesis. A) Timelapse images show HeLa cells treated with anillin RNAi to deplete endogenous anillin, expressing various RNAi-resistant GFP-tagged full-length anillin constructs (green) and mCherry:Tubulin (magenta) as indicated: control, RBD-C2 interface mutant (I/F mutant), NLS mutant, and a combination of the I/F + NLS mutations (I/F + NLS mut.). Times are indicated from anaphase onset. The scale bar is 10 *μ*m. B) A bar graph shows the percentage of cells (Y-axis) that failed ingression (light grey) compared to those that completed ingression (dark grey) for the different constructs in A) as indicated: control (n=15), I/F mutant (n=22), NLS mutant (n=22), I/F + NLS mutant (n=17), and *strong* I/F mutant (Strong I/F mutant; L735D, L736D; n=16). The data were analyzed using Fisher’s exact test (n.s., not significant; *p < 0.05; **p < 0.001; ***p ≤ 0.0001). C) A bar graph shows the time from anaphase onset to the end of ingression (Y-axis) in cells rescued with the various anillin constructs in A): control (n=15), I/F mutant (n=17), NLS mutant (n=14), I/F + NLS mutant (n=5) and Strong I/F mutant (n=6). Bars indicate SD, which are also shown as time on the graph. Data was analyzed and p values were determined by Student’s t test (*p < 0.05; **p < 0.001; ***p ≤ 0.0001). D) A bar graph shows the percentage of binucleate cells (Y-axis) as a readout for cytokinesis failure in populations of asynchronous HeLa cells after anillin RNAi and co-expressing with the indicated RNAi-resistant constructs. Bars show SD (n=3 replicates for each condition). Data was analyzed and p values were determined by the Student’s t test (**p < 0.001; ***p ≤ 0.0001). E) A schematic shows how importin-binding (orange) may increase RhoA-binding (RhoA-GTP in blue; RBD in pink) or reduce its off-rate by stabilizing a favorable conformation of the RBD and C2 domain (green; NLS also shown). The PH domain is in yellow.

To further assess how these mutations affect cytokinesis at the population level, we performed experiments as above on asynchronous populations of HeLa cells. We calculated the proportion of binucleate cells for each construct as a read-out for cytokinesis failure (Figure 5D). There was a significant increase in the proportion of binucleate cells depleted of endogenous anillin and expressing RNAi-resistant I/F mutant anillin (49.5% compared to cells expressing control anillin 23.5%), which was also observed in cells expressing the NLS mutant (47.7%). However, there was an even greater increase in the proportion of binucleate cells expressing anillin with both the NLS and I/F mutations, which was similar to the anillin RNAi control (68.7% vs. 72.3%). This data shows that weakening the interface of the RBD and C2 causes a dramatic change in anillin’s recruitment when importin-binding is lost, suggesting that there is feedback from the C2 domain to the RBD to facilitate RhoA-binding (Figure 5E). To test this hypothesis, we performed *in vitro* experiments to determine how importin-binding influences anillin’s interaction with RhoA. MBP:Anillin RBD + C2 was used to pull down His-tagged RhoA-GDP or RhoA-GTP in the presence or absence of GST:Importin-β (Figure S3). Adding importin-β increased anillin’s affinity for GTP-bound RhoA, suggesting that importin-binding alters the RBD to make it more accessible and/or to stabilize its interaction with active RhoA (Figure 5E).

### Cortical recruitment of anillin relies on the interface between the RBD and C2

Our data suggests that the interface between the RBD and C2 domains contributes to anillin function. While active RhoA-binding makes the C2 more accessible for importin-binding, importin-binding also makes the RBD for accessible for binding to active RhoA. To further test how the interface mediates anillin’s ability to interact with importin-β or RhoA, we created point mutations within the RBD (*strong* I/F mutant; L735D, L736D) to specifically disrupt hydrophobic interactions with the C2 domain (Figure 6A). Based on their position, these mutations should more strongly disrupt the interface vs. the mutations described above (843-DFEINIE to AFAINIA). First, we assessed how the *strong* I/F mutations affect autoinhibition of the NLS in the C2 domain. We previously showed that the C-terminus of anillin fails to localize to the nucleus in interphase cells, but is nuclear after removing the RBD (Beaudet et al., 2017). We measured changes in the nuclear localization of the C-terminus of GFP-tagged anillin with mutations in the NLS, *strong* I/F, or *strong* I/F + NLS. Based on relative levels, their distribution was categorized as nuclear or cytosolic, and only considered if their average fluorescence intensity was over 1500 a.u. As expected, the C-terminus localized primarily to the cytosol in interphase cells (Figure 6B). Introducing the *strong* I/F mutations caused the C-terminus to localize to the nucleus in approximately 64% of the cells, which was lost when the NLS was also mutated (Figure 6B).

**Figure 6.**
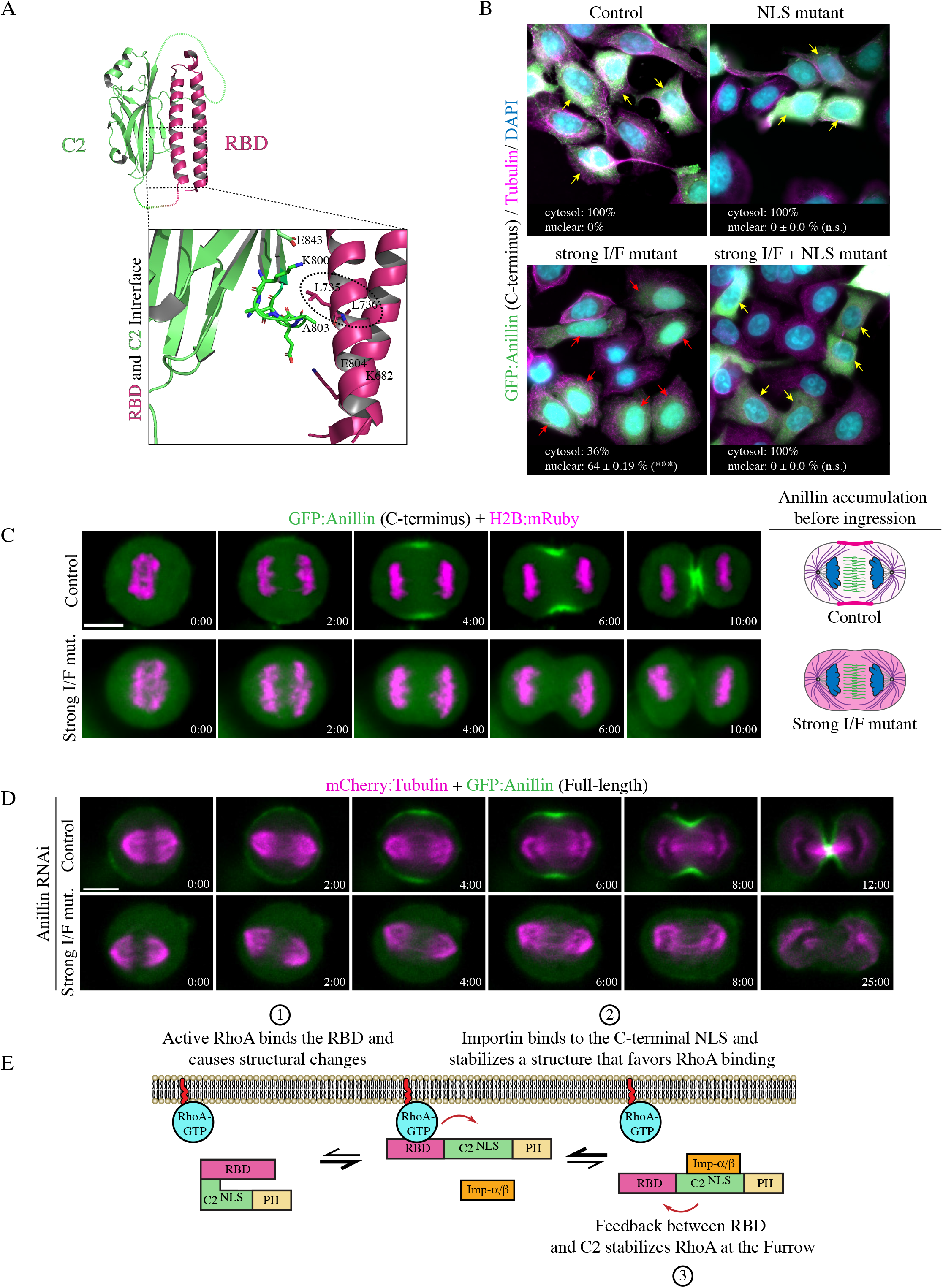
The interface between the RBD and C2 domains is required for feedback for robust cortical recruitment and cytokinesis. A) A ribbon structure shows the intramolecular interface between the RBD (pink) and C2 domain (green). The boxed inset (left) shows the amino acids that form electrostatic and hydrophobic interactions at the interface. The amino acids that were mutated to generate a *strong* I/F mutant are circled (L735D and L736D). B) Images show fixed HeLa cells expressing GFP-tagged anillin C-terminus, or the NLS mutant, *strong* I/F mutant, or *strong* I/F + NLS mutant, stained for GFP (green), tubulin (red), and DAPI (DNA; blue). Yellow arrows point to cytosol localization, while red arrows point to nuclear localization. The percentage of cytosolic vs. nuclear localization is indicated for each condition. SD are shown (n=3 replicates for each condition) and significance was determined using the Student’s t test (n.s., not significant, *** p < 0.0001). C) Timelapse images show HeLa cells expressing GFP-tagged C-terminus anillin or containing the *strong* I/F mutations (green) and H2B:mRuby (magenta; to show chromatin; control, n=18; *strong* I/F mutant, n=7). Times are indicated form anaphase onset. The scale bar is 10 *μ*m. Cartoon cells on the right show changes in the distribution of anillin (pink) at the onset of ingression for control vs. the *strong* I/F mutant. D) Timelapse images show HeLa cells treated with anillin RNAi and coexpressing RNAi-resistant GFP-tagged full-length (control; n=15) or *strong* I/F mutant (n=16) anillin (green), and mCherry:Tubulin (magenta). Times are indicated form anaphase onset. The scale bar is 10 *μ*m. Times are indicated form anaphase onset. The scale bar is 10 *μ*m. E) A cartoon schematic shows how importin-binding enhances the RhoA-mediated recruitment of anillin to the equatorial cortex during cytokinesis. (1) RhoA is activated at anaphase onset and binds to the RBD (pink) to recruit anillin to the equatorial cortex. Active RhoA (blue) increases anillin’s affinity for importin by causing conformational changes between the RBD and C2 domain (green). (2) Near the cortex, importin-β (orange) binds to anillin via the C-terminal NLS, which (3) feeds back by stabilizing a conformation that enhances RhoA binding to the RBD via increasing affinity or reducing dissociation.

Therefore, the *strong* I/F mutations are sufficient to relieve autoinhibition from the RBD, and do not impact importin-binding.

Next, we wanted to compare how the *strong* I/F mutant affects importin-binding after manipulating active RhoA or RhoA-binding as in Figure 3C. We used recombinant MBP:Anillin (RBD + C2) control, RBD mutant, or the *strong* I/F mutant to pull down GFP-tagged Importin-β from lysates +/− active RhoA (Figure 3D). As before, we used over-expression (O/E) of the C-terminus of Ect2 [Myc:Ect2 (C-term)] to generate active RhoA, and Ect2 RNAi to reduce active RhoA. We found that Importin-β:GFP had higher affinity for the *strong* I/F mutant compared to control regardless of the levels of active RhoA, while the RBD mutant showed the opposite trend (Figure 3D). This data supports the idea that the *strong* I/F mutant mimics an ‘open’ conformation of anillin that no longer requires RhoA-binding for increased accessibility of the NLS.

Next, we determined how disrupting the interface affects anillin’s localization and function during cytokinesis. Cells expressing GFP-tagged C-terminus of anillin or with the *strong* I/F mutations, and H2B:mRuby to visualize chromatin, were imaged after anaphase onset. As previously described, anillin localized to the cortex ~2 minutes after anaphase onset and accumulated at the equatorial cortex during ingression. However, anillin containing the *strong* I/F mutations failed to localize to the cortex and remained cytosolic through ingression (Figure 6C). To determine how the *strong* I/F mutant impacts anillin’s function for cytokinesis, RNAi-resistant GFP-tagged full-length anillin or containing the *strong* I/F mutations were imaged in HeLa cells depleted of endogenous anillin and co-expressing mCherry:Tubulin (Figure 6D). Full-length anillin localized similar to the C-terminus as previously described and all cells completed ingression, while the *strong* I/F mutant failed to localize to the cortex and the majority of cells failed to ingress (62.5%; Figure 5B). Cells that successfully ingressed were significantly delayed compared to control cells (18.3 ± 2.4 mins; Figure 5C). Thus, the localization and phenotype of the *strong* I/F mutant resembles constructs containing mutations in the RBD, or in both the I/F (*weak*; 843-DFEINIE to AFAINIA) and NLS. We also assessed how the *strong* I/F mutations impact cytokinesis at the population level and found that the proportion of binucleate cells was similar to the anillin RNAi control (71% vs. 72.3%; Figure 5D). This data suggests that the cortical affinity of anillin is regulated by importin-binding, which we propose feeds back through the interface to influence anillin’s cortical recruitment by RhoA.

## DISCUSSION

Our studies shed light on how anillin’s function for cytokinesis is regulated at the intramolecular level by active RhoA and importin-β. Our previous study showed that cytokinesis is regulated by Ran-GTP, which may coordinate ring positioning with chromatin (Beaudet et al., 21017). We also showed that anillin is a target of this pathway, as it contains a highly conserved NLS that binds to importin-β, and is required for its proper localization and function during cytokinesis (Beaudet et al., 2017). As described earlier, a gradient of active Ran persists around chromatin, and importin-bound cargo form an inverse gradient (Kaláb and Heald, 2008). Having high importins free to bind to cargo near the equatorial cortex could help regulate cortical proteins required for cytokinesis. Here we delve into the mechanism by which importin-binding regulates anillin function. Our model (Figure 6E) is that when the levels of active RhoA increase upon anaphase onset, the binding of active RhoA to the RBD causes a conformational change in anillin that makes the neighbouring C2 domain more accessible to binding partners. One of these partners is Importin-β, which binds to the NLS in the C2 domain. Importin-binding then stabilizes the open conformation to reinforce binding to active RhoA as well as to other factors required for anillin’s recruitment and retention at the equatorial cortex. Our binding data supports this model by showing that active RhoA increases anillin’s affinity for importin, and vice versa. Our localization data also shows the importance of the interface between the RBD and C2 domain, which is required for feedback likely to position the domains for binding RhoA and importin with optimal affinity.

While our work focused on importin-binding, it is important to note that the C2 domain also contains binding sites for microtubules and phospholipids that may be autoinhibited by the RBD. Further, microtubule or lipid binding to the C2 domain also could influence anillin’s affinity for active RhoA. We previously found that there was an inverse correlation with anillin’s localization to microtubules vs. the cortex, depending on the levels of active RhoA or the stability of microtubules (van Oostende Triplet et al., 2014). In addition, a recent study showed that phospholipid binding may facilitate feedback to reinforce RhoA binding (Budnar et al., 2019).

### Importin-binding regulates anillin’s affinity for the equatorial cortex

Anillin is recruited to the equatorial cortex by active RhoA, and is restricted from the polar cortices by astral microtubules. Previous studies showed that reducing active RhoA or stabilizing microtubules with Taxol increase anillin’s localization to microtubules, suggesting that they compete with each other (van Oostende Triplet et al, 2014). We also found that mutating the NLS to reduce importin-binding weakens anillin’s cortical affinity, since its recruitment is delayed and more restricted compared to non-mutant anillin. In addition, disrupting astral microtubules causes the NLS mutant to spread along the cortex, suggesting that the weakened affinity favors a transition in its localization to microtubules (Beaudet et al., 2017). Here we continued to explore how importin-binding affects anillin’s cortical properties by performing FRAP studies. NLS mutations that reduce importin-binding decreased anillin’s mobility and increased its off-rate (Figures 1 and 2). The reduced mobility likely reflects the shift onto microtubules, since we also saw reduced mobility of anillin when microtubules were overextended by MCAK RNAi (Figure 2). Interestingly, when the NLS is mutated, anillin’s affinity for microtubules is not increased per se, since we found via co-sedimentation assays that its affinity is partially reduced (Figure S2). Thus, the reduced mobility via the NLS may be indirect due to an increase in microtubule localization caused by an overall reduced affinity for RhoA. We also saw an increased off-rate with the NLS mutant that we did not see with MCAK RNAi, which may indicate that importin-binding specifically regulates cortical retention.

### RhoA-binding relieves autoinhibition of the C2 domain from the RBD

We previously found that the C2 domain in the C-terminus of anillin had higher affinity for importin and microtubules when the RBD was removed (Beaudet et al., 2017). Since the RBD binds to active RhoA, we proposed a model where active RhoA-binding could induce a conformational change that makes the C2 more accessible. Here, we tested this model by changing the levels of active RhoA, or mutating the RBD and seeing how this influenced importin-binding. In support of our model, when active RhoA levels were high (*e.g*. over-expressed GEF domain of Ect2), we saw increased importin-binding compared to when active RhoA levels were low (*e.g*. Ect2 RNAi), or when the RBD was mutated (Figure 3). Thus, similar to other RhoA effectors, such as mDia, intramolecular autoinhibition could ensure that anillin’s other interactions such as binding to phospholipids, are closely linked to when active RhoA levels increase during mitotic exit, and in the region of the equatorial cortex (Li and Higgs, 2003; Lammers et al., 2005).

### Feedback at the interface between the RBD and C2 domain drives anillin’s equatorial recruitment

Our data shows that the NLS of anillin influences its cortical properties. One hypothesis is that the C2 domain feeds back to the RBD to improve accessibility to active RhoA or reduce its off-rate. Indeed, when the NLS mutations were combined with mutations that weaken the interface between the RBD and C2, anillin failed to be recruited to the cortex and failed to support cytokinesis, similar to mutations that disrupt the RBD (Figures 4 and 5). Introducing mutations predicted to more strongly disrupt the interface similarly failed to localize or function, even though binding to importin increased (Figures 5 and 6). Thus, we hypothesize that communication at this interface enables feedback between the RBD and C2 domain that drives the RhoA-mediated recruitment of anillin. Since the C2 domain has binding sites for other factors, it would be interesting to determine how these other interactions also impact binding to RhoA. For example, as described above, stabilizing microtubules competes anillin from the cortex. Since the microtubule- and importin-binding sites overlap, stabilizing microtubules may outcompete importins and sterically hinder the RBD for optimal RhoA binding. In addition, previous studies suggest that there is cooperativity between anillin’s interaction with phospholipids and RhoA, which could involve a feedback mechanism (Sun et al., 2015; Budnar et al., 2019). The lipid binding site is predicted to be at a loop near the NLS, and lipid binding could increase accessibility of the RBD to RhoA, similar to importin. Anillin likely is also regulated by post-translational modifications that may influence its conformation. For example, a recent study showed that phosphorylation at a site that lies just N-terminal to the RBD enhances anillin’s cortical association and is required for cytokinesis (Kim et al., 2017). This area is under-explored and it would be interesting to see how phosphorylation by different cell cycle kinases influences anillin’s structural changes and function for cytokinesis.

## MATERIALS AND METHODS

### Cell culture

HeLa cells were plated and grown in Dulbecco’s Modified Eagle Medium (DMEM; Wisent), supplemented with 10% fetal bovine serum (FBS; Thermo Scientific), 2 mM L-glutamine (Wisent), 100 U penicillin and 0.1 mg/mL streptomycin (Wisent) and were maintained at 37°C with 5% CO_2_. For transfection, cells were plated in DMEM media without antibiotics (PS), and transfected using Lipofectamine 2000 according to the manufacturer’s protocol (Invitrogen), except that 3 μL of Lipofectamine was used per 2 mL of media with 0.5–2.0 *μ*g DNA and 3 *μ*L of 2 nM siRNAs, as described previously (Yüce et al., 2005; Piekny and Glotzer, 2008). Cells were imaged 24–26 hours after DNA transfection, and 27–30 hours after co-transfection of DNA and siRNAs. Anillin, Ect2 and MCAK siRNAs were used as described previously (Yüce et al., 2005; Piekny and Glotzer, 2008; van Oostende Triplet et al., 2014).

### Plasmids

H2B:mRuby was generated from H2B:GFP, generously provided by G. Hickson (University of Montreal). GFP was replaced with mRuby using *BamHI* (New England Biolabs) and *XbaI* restriction enzymes (New England Biolabs), and the pcDNA3:mRuby2 plasmid was kindly provided by C. Brett (Concordia University). A stable HeLa mCherry:Tubulin cell line was generated previously (van Oostende Triplet et al., 2014). pEGFP-N1:Importin-β was obtained by Addgene, made by Patrizia Lavia (plasmid # 106941). GST:Importin-β was made by cloning Importin-β cDNA from the pEGFP-N1 vector into pGEX-4T using *NcoI* and *NotI* (New England Biolabs). The anillin constructs for mammalian cell expression (GFP-tagged) or protein expression (MBP or GST-tagged) were generated previously (Piekny and Glotzer, 2008; Frenette et al., 2012). The 850 KK 851-DE (NLS mutant), A703E; E721A (RBD mutant; Sun et al., 2015), 837 DFEINIE 843-AFAINIA (*weak* I/F mutant; Piekny and Glotzer, 2008), 735 LL 736-DD (*strong* I/F mutant) and combinations of mutations were generated in the anillin constructs by quickchange PCR. The His:RhoA and Myc:Ect2 (C-terminus) constructs were generated previously (Yüce et al., 2005; Frenette et al., 2012). All constructs were verified by sequencing.

### Microscopy

Cells were fixed for immunofluorescence using 10% trichloroacetic acid (TCA) as described previously (Yüce et al., 2005). Fixed cells were immunostained for microtubules using 1:200 mouse anti-tubulin antibodies (DM1A, Sigma-Aldrich), anillin using 1:200 rabbit polyclonal anti-anillin antibodies (Piekny and Glotzer, 2008), GFP using 1:100 mouse Clones 7.1 and 13.1 (Roche) or 1:200 rabbit anti-GFP polyclonal antibodies generously provided by M. Glotzer (University of Chicago). Anti-rabbit or -mouse Alexa 488 and anti-mouse or -rabbit Alexa 568 (Invitrogen) secondary antibodies were used at a 1:250 dilution. DAPI (Sigma-Aldrich) was added at a 1:1000 dilution (1 mg/mL stock) for 5 minutes before mounting the coverslips onto slides. Fixed cells were imaged using a Leica DMI6000B wide-field microscope with the 63x/1.4 PL APO oil immersion objective (pixel size 0.102 μm), and Z-stacks of 0.5 μm were acquired with a Hamamatsu OrcaR2 camera and Volocity software (PerkinElmer) using a piezo Z stage (MadCityLabs). Image files were exported as TIFFs, which were opened with ImageJ (NIH) and converted into maximum intensity Z-stack projections. Projections and merged colour images were then converted into 8-bit images and imported into Illustrator (Adobe) to make figures.

To perform live imaging, media was replaced with phenol red-free DMEM media. Cells were plated and transfected on 25 mm round coverslips (No. 1.5) placed in a 35 mm Chamlide magnetic chamber (Quorum). Cells were kept at 37°C with 5% CO_2_ using the INU-TiZ-F1 chamber (MadCityLabs). Live imaging was performed on an inverted Nikon Eclipse Ti microscope with a Livescan Swept Field confocal unit (Nikon), using the 60x/1.4 CFI PLAN APO VC oil immersion objective (pixel size 0.27 μm), a piezo Z stage (MadCityLabs), and with the iXON897 EMCCD camera (Andor). Images were acquired with 200 ms exposures using the 488 and 561 nm lasers (100 mW, Agilent) set between 20–40% power, depending on the intensity of fluorescent signals (settings were kept constant for related experiments), and multiple Z-stacks of 0.5 *μ*m were taken every 60 seconds per cell using NIS-Elements acquisition software (Nikon), and a narrow GFP or dual filter (500-544 and 600-665 nm; Chroma). All of the images co-expressing GFP and mRuby probes were spectrally unmixed using the NIS-Elements acquisition software (Nikon). Image files were exported as TIFFs, which were opened with ImageJ (NIH) and converted into maximum intensity Z-stack projections. Projections and merged colour images were then converted into 8-bit images and imported into Illustrator (Adobe) to make figures, or saved as AVI movie files.

### FRAP

To perform FRAP experiments, Cells were plated and transfected on 25 mm round coverslips (No. 1.5) placed in a 35 mm Chamlide magnetic chamber (Quorum). Cells were kept at 37°C with 5% CO_2_ using the INU-TiZ-F1 chamber (MadCityLabs). Live imaging and FRAP was performed using a Nikon C2 laser scanning confocal microscope using the 100X Plan Apo I (NA1.4) objective and Elements 4.0 acquisition software (Nikon). Time lapse images were acquired at resolution of 1024×1024 pixels every 2 seconds for a total of 60 seconds. Two time points were acquired prior to photobleaching and for up to 1 minute after photobleaching. Photobleaching of control and experimental ROI’s took place for a total of 5 seconds by pulsing 10 times with the laser power set between 20–50%. Movies were exported as TIFFs then opened with ImageJ (NIH) for analysis. Fluorescence intensity of bleached and non-bleached ROIs were corrected for acquisition bleaching and normalized to background. The average intensity of ROIs was plotted over time using Prism software and the signal recovery rate and maximum recovery of signal was determined using non-linear regression fitting with the following formula:

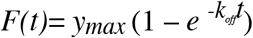

Where *F(t)* describes the fraction of fluorescence recovery for each time point after photobleaching compared to the pre-bleached signal, *y_max_* describes the maximum fluorescence recovery after photobleaching, and *k_off_* describes the dissociation rate. The half-life (τ_1/2_) or time taken to reach half the maximum fluorescence recovery after photobleaching was determined by the following equation:

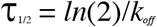

And the immobile fraction was determined as follows:

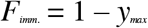

Where, *F_imm_* is the percent difference between the initial fluorescence signal before photobleaching and the maximum signal recovered (*y_max_*) after photobleaching.

### Protein purification, pull downs, and PIP strips

The following proteins were made from *E. coli* BL21 cells: His:RhoA, GST, GST:Importin-β, GST:Anillin (AHD; 608-940), MBP, MBP:Anillin (C2; 750-872), MBP:Anillin (RBD + C2; 672-872), and MBP:Anillin (C-term; 608-1087), as well as containing mutations in the NLS (850 KK 851-DE), RBD (A703E; E721A), or interface (L735D; L736D). Bacteria were resuspended in lysis buffer [2.5 mM MgCl_2_, 50 mM Tris, 150 mM NaCl pH 7.5, 0.5% Triton X-100, 1 mM dithiothreitol (DTT), 1 mM phenylmethanesulfonyl fluoride (PMSF) and 1 X protease inhibitors (Roche)], incubated with 1 mg/mL lysozyme on ice for 30 minutes, then sonicated three times. Extracts were incubated with pre-equilibrated amylose resin (New England Biolabs) or glutathione sepharose 4B (GE Lifesciences) for 5 hours or overnight at 4°C with rotation. After washing, beads were stored as a 50% slurry at 4°C or eluted in equivalent volumes of 100 mM maltose or 10 mM glutathione (pH 8.0) on ice for 2 hours. Protein concentration was determined by running samples by SDS-PAGE and measuring the density of bands in comparison to known concentrations of BSA and/or by Bradford assay for eluted proteins.

To test for binding, proteins were pulled down from cell lysates after transfection or using eluted recombinant purified protein. Transfected Hela cells were lysed in 50 mM Tris pH7.6, 150 mM NaCl, 5 mM MgCl_2_, 0.5% Triton X-100, 1 mM DTT, 1 mM PMSF with 1 X protease inhibitors (Roche) and incubated with 5-10 *μ*g of purified MBP-tagged anillin or GST-tagged importin-β protein on beads at 4°C overnight. Eluted proteins were added to beads at final concentrations as indicated. After binding, beads were washed 3–4 X with 50 mM Tris pH 7.6, 150 mM NaCl, 5 mM MgCl_2_ before adding SDS sample buffer to denature the proteins for SDS-PAGE. All samples were run by SDS-PAGE and wet-transferred to nitrocellulose membrane for western blotting. All blots were reversibly stained with Ponceau S to show total protein. The blots were blocked with 5% milk for 20 minutes, then incubated with either mouse anti-MBP antibodies at a dilution of 1:5000 (New England Biolabs) or 1:2500 mouse anti-GFP antibodies (Roche) in 1× PBS-T (0.140 M NaCl, 2.7 mM KCl, 10 mM Na_2_HPO_4_, 1.8 mM KH_2_PO_4_, 0.5% Triton X-100) for 1-2 hours at room temperature. After washing the membrane 3-4 X with 1 X PBS-T, secondary antibodies [anti-rabbit-HRP or anti-mouse-HRP (Cedarlane)] were added as per manufacturer’s instructions in 1× PBS-T for 1 hour. The blots were developed using enhanced chemiluminescence (ECL) western blotting detection reagents (GE Amersham) and visualized on a GE Amersham Imager 600.

PIP strips (Echelon Biosciences) were blocked with 3% BSA in 1× PBS-T at 4°C overnight. PIP strips were removed from blocking solution and incubated with 5 μg/mL of purified recombinant protein for 1 hour at room temperature. After washing 3–4 X with 1× PBS-T the strips were incubated with 1:5000 mouse anti-GST antibodies (Sigma). The secondary antibodies (anti-rabbit-HRP or anti-mouse-HRP) were then added in 1× PBS-T for 1 hour. After washing the membranes 3-4 X with 1 X PBS-T, the signal was developed using enhanced chemiluminescence (ECL) western blotting detection reagents (GE Amersham) and visualised on a GE Amersham Imager 600. Images were converted to 8-bit by ImageJ, and made into figures using Adobe Photoshop and Illustrator (Adobe).

### Co-sedimentation assays

Microtubules were prepared from lyophilized microtubules (Cytoskeleton) as per manufacturer’s instructions in resuspension buffer (15 mM PIPES, 1 mM MgCl_2_, 20 *μ*M Taxol; Bioshop) at room temperature for 10–15 minutes with gentle mixing. Aliquots of 45.5 *μ*M were flash frozen and stored at −80°C, then thawed in a circulating water bath and diluted with resuspension buffer to 9.1 *μ*M. Purified anillin and importin proteins were pre-spun by centrifugation at 279,000 g for 30 minutes at room temperature. Co-sedimentation reactions were prepared in 200 μL polycarbonate tubes (Beckman Coulter) with 0.5–4 μM of microtubules and 1 μM of purified anillin, or 4 μM of microtubules, 1 μM of purified anillin and 0.2–1.5 μM of purified importin-β, 150 mM NaCl and BRB80 buffer (80 mM PIPES pH 6.8, 1 mM EGTA, 1 mM MgCl_2_) containing 1 mM DTT and 10 μM Taxol. The reactions were kept at room temperature for 15 minutes, then centrifuged at 279,000 g for 30 minutes. The supernatants were collected, and pellets were washed and re-suspended in BRB80. Sample buffer was added to supernatants and resuspended pellets, which were then run by SDS-PAGE and stained with Coomassie Blue. Scanned gels were analyzed in ImageJ to measure the concentration of proteins using line plots to determine pixel intensities that were imported into Excel (Microsoft). After correcting for background, the average intensity was determined. The average amounts of bound anillin (Y axis) were plotted against free microtubule concentration (X axis) using GraFit version 7.0.3. The K_D_ and binding capacity were determined using non-linear regression (n=3 assays). For the competition assay, after correcting for background, the pixel intensity of anillin in the supernatant was divided by the pixel intensity of anillin in the pellet to obtain a ratio of anillin (S/P) for each concentration of importin-β. Anillin ratios for importin-β concentrations 0.2-1.5 μM were compared to control and data was analyzed using the Student’s t-test (n=3 assays).

### Quantification

The breadth and ratio of cortical vs. cytosol accumulation for anillin were performed using ImageJ. Maximum intensity Z-projections were generated for each cell, and the breadth was determined using a line scan drawn along the cell cortex. The breadth was determined as the number of pixels above half of the maximum intensity (width of the peak) after subtracting cytosol levels, which was divided by the total number of pixels to give a ratio of breadth to length. To measure the ratio of cortical accumulation vs. cytosol, the average intensity was determined in an area drawn around the cortex from one pole to the other. This value was then divided by the average fluorescence intensity from the average of 3 ROI’s in the cytosol. Similar timepoints were selected for anaphase just before ingression based on time from anaphase onset. All data was imported into Excel (Microsoft), where calculations were performed including standard deviations and student t tests, and to generate graphs. All of the images and graphs were transferred to Illustrator (Adobe) to make figures.

## Author Contributions

A.P. and D.B. wrote the original manuscript and conceptualized and designed the methodology for experiments. D.B., N.P. and N.S. performed the experiments and analyzed the data. D.B. made the data into figures for visualization. Funds were acquired by A.P. Supervision was carried out by A.P. and D.B.

## Acknowledgements

We thank C. Law for help with imaging studies as part of the Centre for Microscopy and Cellular Imaging at Concordia University. We thank C. Brett (Concordia University), I. Cheeseman (Whitehead MIT) and G. Hickson (University of Montreal) for reagents.

---

**Figure S1. Related to Figure 1. Anillin has different cortical properties during early vs. late ingression** A) Cartoon cells show how the stage of ingression was determined by the ratio of the length of the ingressed cortex (B) over the width of the cell at the equator (A); where *R* > 0.6 was ‘early’ and *R* < 0.6 was ‘late’. B) Timelapse images show FRAP of HeLa cells expressing mCherry:Tubulin (magenta) and GFP:Anillin (full-length; green) during early or late ingression. The boxed insets show the ROI’s that were photobleached in Fire LUTs. The scale bar is 10 μm. Indicated times are before (−) or after (+) photobleaching. C) A graph shows the fraction of fluorescence recovery (Y - axis) over time (X-axis, seconds) for anillin during early (n=31) and late (n=15) ingression. Bars show standard deviation (SD). The table shows the maximum recovery (*y_max_*), dissociation rate (*k_off_*) and half-life (τ_1/2_) of anillin during early and late ingression. Standard errors are shown in parentheses (SEM).

**Figure S2. Related to Figure 2. The microtubule-binding region partially overlaps with the C-terminal NLS** A) Images show Coomassie-stained gels of co-sedimentation assays using 1.5 μM purified microtubules (MTs) with 1.5 μM MBP-tagged anillin C2 (top) or NLS mutant (bottom; S, supernatants; P, pellets). The table shows their mean binding capacity and dissociation coefficients, along with the SD. The graph shows bound anillin C2 (μM; Y-axis) plotted against free microtubules (μM; X-axis). Bars show SD (n=3 replicates). B) Images show Coomassie-stained gels of co-sedimentation assays using 1.5 μM purified MTs incubated with 1.5 μM MBP-tagged anillin C2 and 0-1.5 μM GST-tagged importin-β (S, supernatants; P, pellets). The bar graph shows the ratio of anillin (C2) in the supernatant (S) vs. bound (pellet; P) for the indicated concentrations (μM) of GST:Importin-β. Bars show SD (n=3 replicates). Data was analyzed and p values were determined by Student’s t test (n.s., not significant; *p < 0.05). C) Immunoblots show PIP (phospholipids) strips incubated with purified GST, GST-tagged anillin (AHD) control or NLS mutant protein. The schematic shows the different lipids on the blots.

**Figure S3. (Related to Figure 4 and 5) Importin-binding enhances anillin’s affinity for active RhoA** An immunoblot shows pull downs of purified recombinant His-tagged RhoA pre-loaded with GDP or GTP with MBP-tagged anillin (RBD + C2) +/− purified GST-tagged importin-β as indicated. Inputs are on the left.

