## Supplemental Figure 1 for "Intramolecular regulation of anillin during cytokinesis"

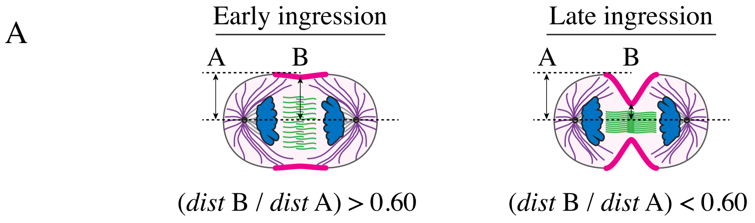


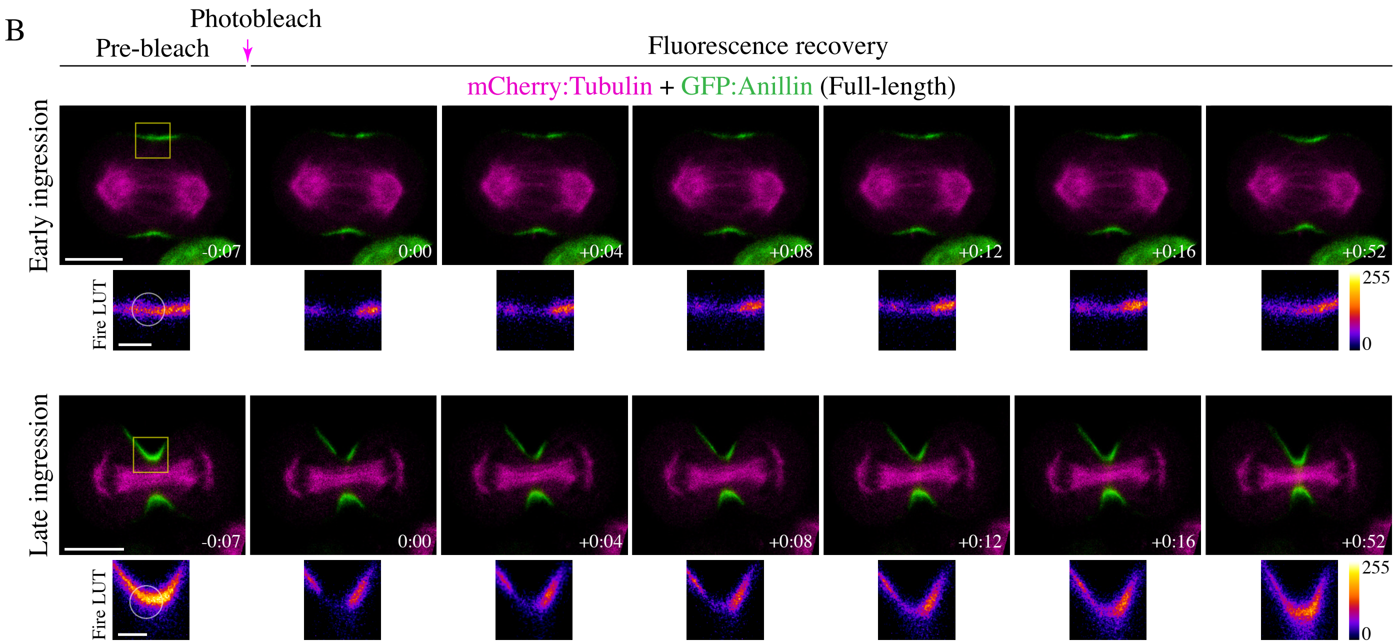


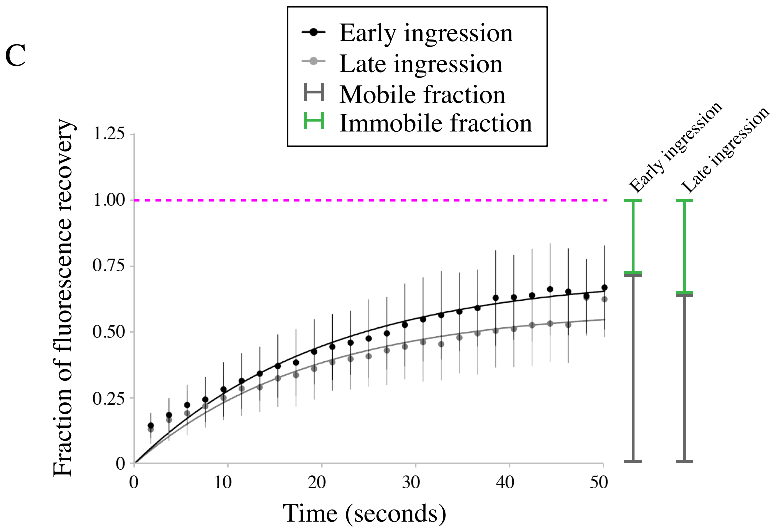


| Anillin's cortical affinity parameters | Early ingression | Late ingression |
| --- | --- | --- |
| $y_{max}$ | 0.7233 (0.02110) | 0.5953 (0.02423) |
| $k_{off}$ ($s^{-1}$) | 0.04717 (0.003119) | 0.05094 (0.004947) |
| $\tau_{1/2}$ (s) | 14.70 | 13.61 |

**Figure S1**
