## Supplementary figures and images for "Intramolecular regulation of anillin during cytokinesis"

### Supplemental Figure 2

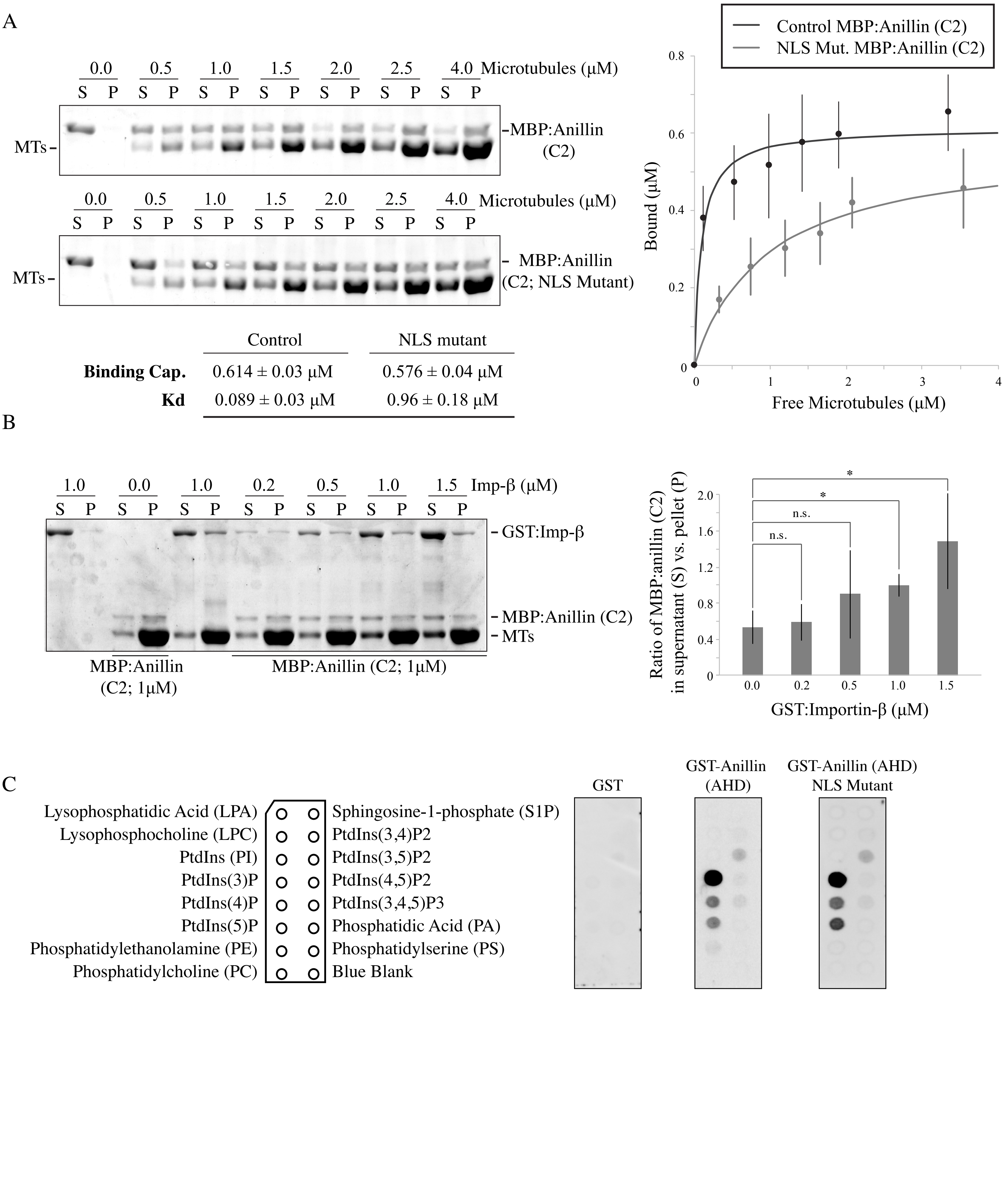

### Supplemental Figure 3

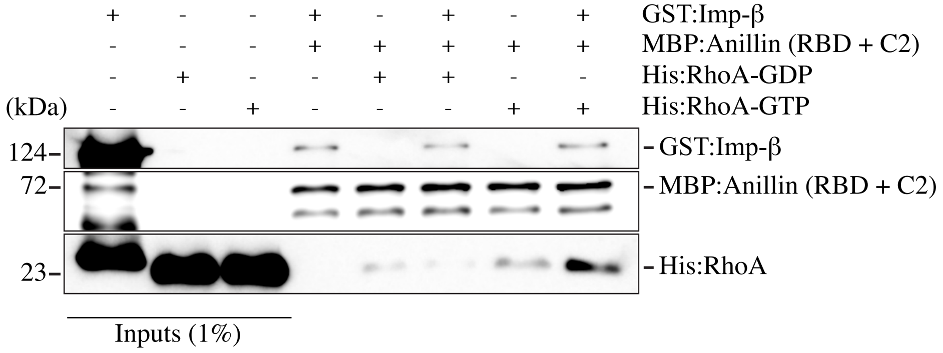
